# Effect of hydroxyapatite microspheres, amoxicillin-hydroxyapatite and collagen-hydroxyapatite composites on human dental pulp-derived mesenchymal stem cells

**DOI:** 10.1101/2021.01.22.427754

**Authors:** Yasmine Mendes Pupo, Lidiane Maria Boldrini Leite, Alexandra Cristina Senegaglia, Lisiane Antunes, Jessica Mendes Nadal, Eliane Leal de Lara, Rafael Eiji Saito, Sandra Regina Masetto Antunes, Paulo Vitor Farago

**Author notes:** Correspondence: Department of Restorative Dentistry, Postgraduate Program in Dentistry, Federal University of Parana, Av. Prefeito Lothário Meissner 632, Jardim Botânico, Curitiba/PR, Brazil, CEP 80210-170.

## Abstract

The aim of this study was to evaluate the *in vitro* behavior of human dental pulp mesenchymal stem cells (hDPSCs) cultured on scaffolds of three hydroxyapatite-based materials: hydroxyapatite microspheres [HAp]; amoxicillin-hydroxyapatite composite [Amx-HAp]; and collagen-hydroxyapatite composite [Col-HAp]. These hydroxyapatites (HAps) were synthesized through three methods: microwave hydrothermal, hydrothermal reactor (teflon pouches), and precipitation, respectively. We performed an *in vitro* experimental study using dental pulp stem cells obtained from samples of third molars and characterized by immunophenotypic analysis. Cells were cultured on scaffolds with osteogenic differentiation medium and were maintained for 21 days. Cytotoxicity analysis and migration assay of hDPSCs were evaluated. Each experiment was performed in triplicate. Data analysis was performed using Kruskal-Wallis test and Dunn’s post-hoc test. After 21 days of induction, no differences in genes expression were observed. hDPSCs highly expressed the collagen IA and the osteonectin at the mRNA, which indicated these genes plays an important role in odontogenesis regardless of induction stimulus. Cytotoxicity assay using hDPSCs demonstrated that Col-HAp group presented a number of non-viable cells statistically lower than the control group (p=0.03). In the migration assay after 24h, biomaterials HAp, Amx-HAp, and Col-HAp revealed the same migration behavior for hDPSCs observed to the positive control. Col-HAp also provided a statistically significant higher migration of hDPSCs than HAp (p=0.02). The migration results in 48h for HAp, Amx-HAp, and Col-HAp was intermediate from those achieved by control groups. There was no statistical difference between positive control and Col-HAp (p>0.05). In general, Col-HAp scaffold showed better features for these dynamic parameters of cell viability and cell migration capacities for hDPSCs, leading to suitable adhesion, proliferation, and differentiation of this osteogenic lineage. These data present high clinical importance because Col-HAp can be used in a wide variety of therapeutic areas, including ridge preservation, minor bone augmentation, and periodontal regeneration.

## 1. Introduction

The developing of new biomaterials or their structural changing has been intensively proposed in order to enhance the final properties of novel biomedical devices.[1] Hydroxyapatite [HAp (Ca_10_(PO_4_)_6_(OH)_2_)] is thermodynamically stable in body fluid due to its crystalline state and has a very similar composition to mineral bone.[2] Thus, numerous techniques to synthesize hydroxyapatite-based materials at elevated temperatures (750–1000 °C) have been used in order to achieve accurate control of their structure. These procedures may be divided into two major groups: wet-chemical methods and solid-state reactions.[3,4] However, other authors have been reported a more detailed classification as: 1) dry methods involving solid-state and mechanomechanical ones; 2) wet methods involving chemical precipitation, hydrolysis, sol-gel, hydrothermal, emulsion and sonochemical ones; 3) high-temperature processes including combustion and pyrolysis method; 4) synthesis based on biogenic sources; and 5) combination procedures.[2]

The method chosen has a remarkable influence on the morphology of the HAp obtained and can modify its crystallographic and chemical structure of powder.[2] In addition, preparation of nanosized HAp is also related to a number of other problems, including difficulties in controlling geometry, size and size distribution, crystallinity, stoichiometry, and degree of particle agglomeration.[2] It is well known that *in vitro* and *in vivo* biological and mechanical properties of HAp are strongly affected by its structural characteristics; hence, extensive efforts have been devoted to engineer the HAp crystals, in particular, by developing new routes or modifications of pre-existing methods.[2]

In that sense, the preparation of modified HAps or its composites requires a careful biological investigation to guarantee the safe use. Cellular studies involving human dental pulp stem cells (hDPSCs) have been widely explored as an prior parameter to predict safety of dental material. These hDPSCs can be easily isolated from dental medical wastes, extracted teeth, and expanded *ex vivo*. hDPSCs are a kind of mesenchymal stem cell with the potential for cell-mediated therapy and tissue engineering applications because of their low morbidity after collection, easy surgical access, ability to be cryopreserved, ability to be recombined with many scaffolds and immune-privilege and anti-inflammatory abilities.[5–7] However, bone formation from hDPSCs requires a special structure provided by scaffolds which should provide an appropriate environment for cellular attachment, growth, and differentiation.[8] In that sense, novel hydroxyapatite-based materials need to be tested using hDPSCs in order to investigate their potential as a suitable environment for cell growth and that at the same time provide safety for clinical use.

In this study, the following HAP-based materials were obtained and characterized: hydroxyapatite microspheres [HAp]; amoxicillin-hydroxyapatite composite [Amx-HAp]; and collagen-hydroxyapatite composite [Col-HAp]. These scaffolds could play a critical role in attachment, survival, migration, proliferation, and differentiation. However their effect on hDPSCs remains unknown. Therefore, we investigated the effects of HAP-based materials by *in vitro* cell culture-based assays on osteogenic differentiation, cytotoxicity, and migration of hDPSCs. Our null hypothesis was that these HAp-based materials, in particular Amx-HAp and Col-HAp, reduce osteogenic differentiation, cause high hDPSC death and prevents cell migration process.

## 2. Materials and Methods

This work was approved by the Research Ethics Committee of the Federal University of Paraná-Brazil (CAAE: 65794316.4.0000.0102). All samples were collected after completing the informed consent form. Three HAP-based materials were tested: hydroxyapatite nanosheet-assembled microspheres [HAp]; amoxicillin-hydroxyapatite composite [Amx-HAp]; collagen-hydroxyapatite composite [Col-HAp]. The tests were performed with hDPSCs from three different donors.

### 2.1 Synthesis and characterization of HAp-based materials

#### 2.1.1 Synthesis of HAp-based materials

HAp, Amx-HAp, and Col-HAp were synthesized through three synthesis methods: microwave-hydrothermal, reactor-hydrothermal (teflon pouches), and precipitation, respectively.

Two aqueous solutions of 0.0835 mol/L calcium acetate and 0.0501 mol/L ammonium dihydrogen phosphate were prepared as precursors to obtain hydroxyapatite microspheres (HAp) by the microwave-hydrothermal method at 1.67 stoichiometric ratio of Ca/P. The complexing agent citric acid monohydrate was added after mixing the calcium and phosphorus precursors until a pH = 4 was achieved. Urea was then added at 0.016 mol/L concentration. The reaction mixture was transferred to reactors that were inserted into a microwave oven (Miliestone STAR D). The microwave oven was set for a gradual heating from room temperature (20°C) to 180°C with at 1000 W power. The synthesis time for reaching 180°C was 5 min. The precipitate vacuum filtered, washed twice, and oven dried.

The amoxicillin-hydroxyapatite composite (Amx-HAp) was prepared by the reactor-hydrothermal method using teflon pouches.[9] Aqueous solutions of calcium acetate and ammonium dihydrogen phosphate at the same concentrations (0.0835 and 0.0501 mol/L, respectively) were used. Citric acid monohydrate was added to calcium acetate solution under magnetic stirring for 30 min to adjust pH at 4. This final solution was dripped under the ammonium dihydrogen phosphate solution and urea at 0.016 mol/L was added. The synthesis procedure was carried out into an autoclavable reactor at 180°C for 24 h. After drying, amoxicillin was blended by milling at a final ratio of 4.80 mg HAp:1 mg Amx. This method was performed into a high-density polypropylene flask using yttria stabilized zirconia spheres in ethanol as grinding agent for 3 h. After milling, Amx-HAp was oven dried for 24 h at 35°C.

The collagen-hydroxyapatite composite (Col-HAp) was obtained by the precipitation method. This reaction was performed by adding 1.2 mol/L phosphoric acid (85% pure) into an aqueous suspension of 2.0 mol/L calcium hydroxide at a Ca/P molar ratio of 1.67 under magnetic stirring at 40°C. The phosphoric acid dropping was controlled at 1 drop/s. The pH was then adjusted to 10.0 using ammonium hydroxide (28% pure). The precipitate was aged for 36 h, vacuum filtered and oven dried. Collagen was also blended by ball milling at a ratio of 4.80 mg HAp:1 mg Col.

#### 2.1.2 Characterization of HAp-based materials

Hydroxyapatite-based materials were characterized by Fourier-transformed infrared (FTIR) spectra that were recorded from 4000 to 400 cm^−1^ on a Biorad Excalibur Series (FTS-3500 GX) IR spectrophotometer using KBr pellets with 32 scans and resolution of 2 cm^−1^.[10] Morphological characterization was performed using field-effect emission gun scanning electron microscopy (FEG-SEM, TESCAN, model MIRA 3, Brno, Czech Republic) at various magnifications. The samples were previously deposited on polished stubs and then sputter-coated with gold. The average particle size of biomaterials were obtained by photon correlation spectroscopy (Zetasizer Nanoseries, Malvern Instruments, Malvern, UK) after diluting each sample in ultrapure water (1:500, v/v) with no previous filtration and sonicating for 30 min.

### 2.2 Collection and characterization of human dental pulp-derived mesenchymal stem cells (hDPSCs)

Previously to collect permanent teeth, the dental surgeon requested to the patient perform a buccal rinse with chlorhexidine to remove possible contaminants, guaranteeing the integrity of the material collected in the Dentistry Department of the Federal University of Paraná (Department of Stomatology). The tooth (3rd molar) was withdrawn from the collection flask and washed in phosphate buffer saline (PBS) (Gibco™ Invitrogen, NY, USA). In petri dishes, the tooth pulp fragments were mechanically removed using endodontic file (Hedstroem - type H). In the sequence cell suspension was dissociated by collagenase type II (Gibco™, Carlsbad, USA) under agitation at 37 °C for 1 hour, filtered (40μm), diluted in PBS and centrifuged. Subsequently the pulp cells were plated in a flask with 25 cm^2^ growth area containing 5 ml of Iscove’s Modified Dulbecco’s medium (IMDM) (Gibco™, Carlsbad, USA) supplemented with 20% fetal bovine serum (FBS) (Gibco™, Carlsbad, USA) and 1% antibiotic (100 units/mL penicillin and 100 μg/mL streptomycin) (Gibco™, Carlsbad, USA) and were stored in an incubator at 37°C with 5% CO_2_ tension. The culture medium was replaced three times a week. When the cultures reached about 80-90% confluence, enzymatic dissociation was performed using trypsin/EDTA (0.25%) (Gibco™, Carlsbad, USA). Inactivation of the enzyme was performed with FBS and IMDM. The cell suspension was centrifuged, the supernatant discarded and the cells counted and plated again. All experiments were performed between the third (P3) and fifth passage (P5) of the cells.

#### 2.2.1 Characterization of human dental pulp-derived mesenchymal stem cells (hDPSCs)

Immunophenotypic analysis was performed by staining 5 × 10^5^ expanded hDPSCs. The cells were incubated with conjugated monoclonal antibodies against the following antigens: CD14 (fluorescein isothiocyanate (FITC) conjugated), CD45 (FITC conjugated), CD19 (FITC conjugated) CD90 (phycoerythrin (PE) conjugated), CD73 (PE conjugated), HLA-DR (peridinin chlorophyll protein (PerCP), CD34 (allophycocyanin (APC) conjugated, CD105 (APC conjugated) and CD29 (APC conjugated). All antibodies are from BD Pharmigen™ (San Jose, CA, USA) and they were used as suggested by the manufacturer. The incubations were performed at room temperature for 30 min. Isotype identical antibodies served as controls. After incubation, the cells were washed with PBS and fixed with PBS containing 1% paraformaldehyde (Sigma-Aldrich, São Paulo, Brazil). Samples were acquired (100,000 cells) in BD FACSCalibur flow Cytometer (Becton Dickinson, San Jose, USA) and quantitative analyses were performed by FlowJo software v8.0.2 (Tree Star, Ashland, USA).

### 2.3 *In vitro* cell culture – based assays

#### 2.3.1 Osteogenic Differentiation

Positive control of reactions was performed by inducing the differentiation of hDPSCs using an osteogenic induction medium (Differentiation Basal Medium-Osteogenic, Lonza, Walkersville, MD, USA) 37°C with 5% CO_2_ tension and were maintained for 21 days. The culture medium was replaced three times every seven days. The differentiation was under the same conditions for each biomaterial tested: HAp, Amx-HAP and Col-HAp. Osteoblastic differentiation was confirmed by mineral deposition of the culture, which was assessed by Alizarin red S (Sigma-Aldrich, São Paulo, Brazil) staining using an optical microscope (NIKON Eclipse Ni, Tokyo, Japan).

After 21 days total RNA was extracted with the TRIzol Reagent (Invitrogen, NY, USA), according to the manufacturer’s instructions. To degrade any contamination of DNA, RNA was treated with DNAse I (Qiagen, Germantown, USA). Complementary DNA (cDNA) was synthesized from 1 μg of total RNA, using oligo-dT primers (USB Corporation, Cleveland, USA) and a reverse transcriptase kit (ImPROm-II, Promega, USA), according to the manufacturer’s instructions. Polymerase chain reaction (PCR) was carried out with 100 ng of cDNA as the template, 5 pmol of each primer, Taq polymerase and reaction mix (IBMP, Brazil). Primers used were as follows: COL1A, 5’GGCCATCCAGCTGACCTTCC3 and; 5’CGTGCAGCCATCGACAGTGAC3’ OSTEONECTINA, 5’ACATCGGGCCTTGCAAATACATCC 3’ and 5’ GAAGCAGCCGGCCCACTCATC 3’. We subject 20 μl of RT-PCR products to electrophoresis in a 2% agarose gel. The bands obtained were visualized by GelRed^®^ Nucleic Acid Gel Stain and photographed under ultra-violet transillumination (UV White Darkroom, UVP Biomaging Systems, USA). Glyceraldehyde-3-phosphate dehydrogenase (GAPDH) transcript was used as internal control (5’ GGCGATGCTGGCGCTGAGTAC 3’ and 5’ TGGTTCACACCCATGACGA 3’).

#### 2.3.2 Cytotoxicity Analysis

One hundred thousand hDPSCs were plated in two T25 culture flasks and cultured in incubator at 37°C with 5% CO_2_ tension and 95% humidity for five days with IMDM medium and 10% FBS. The differentiation was realized for each condition tested: HAp, Amx-HAP, Col-HAp, and control groups. The evaluation was performed using 7-aminoactinomycin D (7-AAD) (BD Pharmingen, San Jose, USA) is a fluorescent dye with high affinity for DNA, which is used as a cell viability dye. Cells with compromised membranes will be stained with 7-AAD, evidencing non-viable cell. Shortly after cellular dissociation, 7-AAD dye was added to the cells and then incubated for 30 minutes at room temperature (~22°C). Samples were acquired in BD FACSCalibur flow cytometer (Becton Dickinson, San Jose, USA) and analyzed by Software FlowJo Version 8.0.2 (Tree Star, Ashland, USA).

#### 2.3.3 Migration Assay

To test whether biomaterials stimulate the chemotaxis of hDPSCs, the cell migration assay was performed using the Boyden chamber with 8.4 mm diameter and 8 μm pore size (ThinCert ™, Greiner Bio-One, Americana, Brazil) in a 24-well plate (CELLSTAR^®^, Greiner Bio-One, Americana, Brazil). The method used for this assay was adapted from Galler *et al*. [11] Duplicate experiments were performed with the three hDPSCs samples. Prior to the start of the experiment, hDPSCs were cultured for four days with 1% FBS. The total of 2 x 10^5^ HDPSCS in IMDM with 1% FBS was seeded at the top of the boyden chamber in the volume of 50 μL. After four hours, 150 μL of the same solution was added to the upper chamber and the bottom was filled with each biomaterial in IMDM with 10% PBS, IMDM with 1% FBS (negative control) or IMDM with 20% PBS (positive control) in the final volume of 400 μL. The experiment was incubated at 37°C and 5% CO_2_ for 24h and 48h. The cells of the upper chamber were carefully removed with a cotton swab. The cells that migrated to the lower chamber were washed with 70% ethanol (Labmaster, Pinhais, Brazil) and incubated with 0.2% crystal violet (Sigma-Aldrich, São Paulo, Brazil) for 10 min. The excess dye was removed with distilled water and the samples were transferred to a 96-well plate (CELLSTAR^®^, Greiner Bio-One, Americana, Brazil). The assay was evaluated on a microplate reader in the 595 nm filter (Nova Analítica, São Paulo, Brazil).

### 2.4 Statistical Analysis

The normal distribution of the migration assay was assessed by the D’Agostino and Pearson test. The data were classified as non-parametric. The comparison between each biomaterial with the negative and positive controls, as well as the biomaterials between them, were analysed with the Kruskal-Wallis test with Dunn post-test. The results were presented in mean absorbance ± SD. Results were considered significant at p <0.05. The analysis were performed using GraphPad Prism version 5.03 for Windows (GraphPad Software, San Diego, USA).

## 3 RESULTS

### 3.1 Synthesis and characterization of HAp-based materials

HAp-based materials were successfully obtained by three proposed methods and their characterization data are depicted in Fig 1. Considering the X-ray diffraction (XRD) analysis, the XRD patterns of HAp (Fig 1A), Amx-HAp (Fig 1B), and Col-HAp (Fig 1C) presented the typical crystalline peaks achieved for pure HAp (JCPDS n° 09-0432), in which the three most intense peaks were assigned at 2θ of 31.8°, 32.2°, and 32.9° and were corresponded to the planes (211), (112), and (300), respectively. The hydroxyapatite microspheres (HAp) were obtained by the microwave-hydrothermal method and displayed wide peaks, which can be associated with the low degree of crystallinity due to the short crystallization time.[12] Amx-HAp showed more defined and higher intensity crystalline peaks. Col-HAp demonstrated an intermediate pattern of crystallinity.

**Fig 1.**
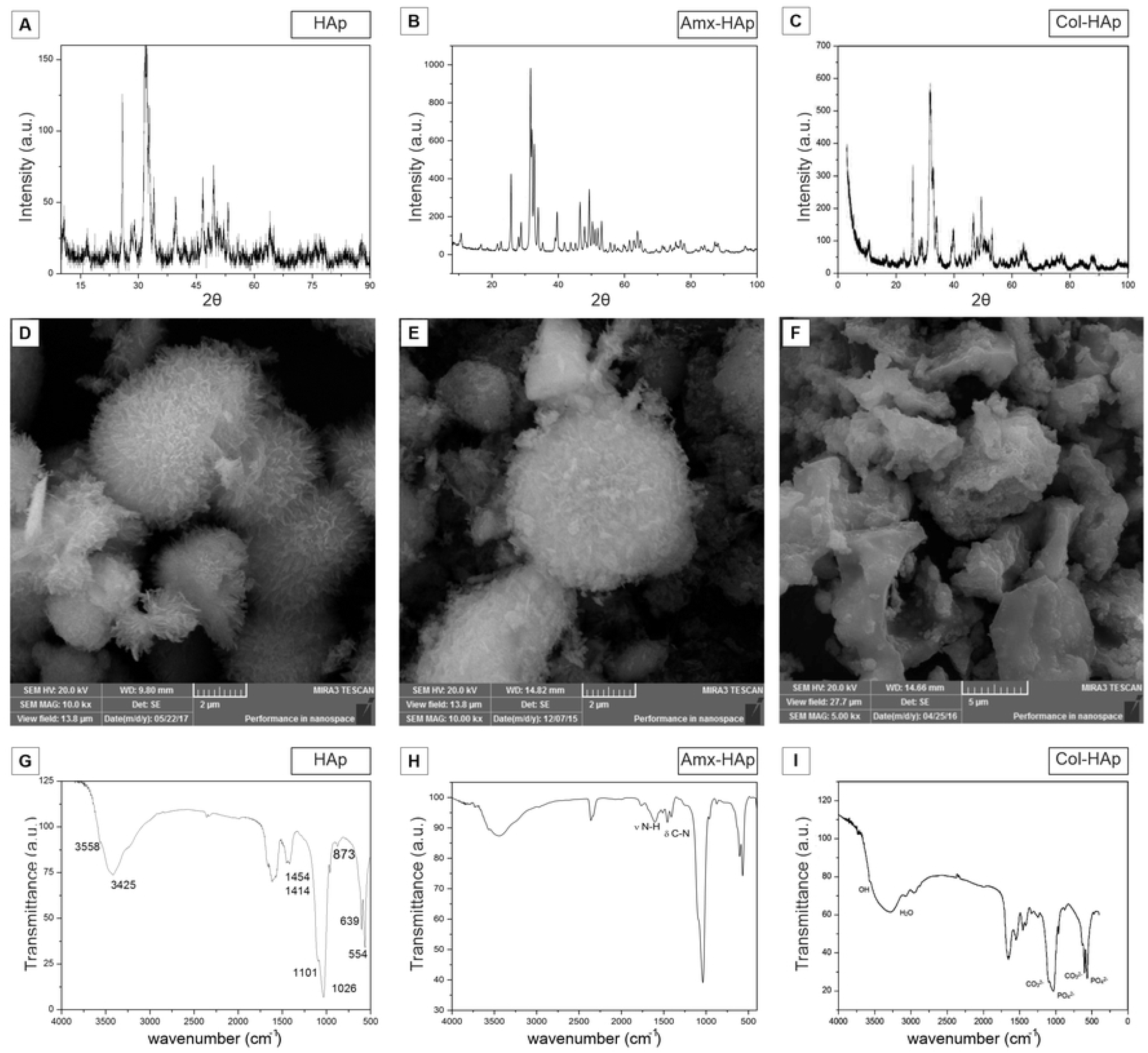
Characterization of HAp-based materials: [HAp: hydroxyapatite nanosheet-assembled microspheres; Amx-HAp: amoxicillin-hydroxyapatite composite; Col-HAp: collagen-hydroxyapatite composite;]: A-C) X-ray diffractogram; D-F) SEM-FEG photomicrographs (SEM HAp and Amx-HAp: 10 kx magnification; SEM Col-HAp: 500× magnification). G-I) FTIR spectra.

The microwave-hydrothermal method provided a biomaterial (HAp, Fig 1D) as nanosheet-assembled microspheres. These microparticles presented a mean particles size of 2305 ± 532 nm. The Amx-HAp composite (Fig 1E) showed similar morphological features to HAp with a mean particles size of 2743 ± 538 nm. The Col-HAp composite (Fig 1F) exhibited irregular microparticles with smooth surface and mean particles size of 7814 ± 653 nm.

HAp synthesized by the microwave-hydrothermal procedure (Fig 1G) showed broad bands at 3425 and 1620 cm^−1^ were attributed to adsorbed water, while the sharp peak at 3558 cm^−1^ was assigned to the stretching vibration of the lattice OH^−^ ions and the medium sharp peak at 639 cm^−1^ was assigned to the OH deformation mode. Some typical bands for PO_4_^3−^ were observed at 554, 873, 1026, and 1101 cm^−1^. These bands confirmed that this biomaterial was consistent to HAp in accordance to literature.[13]

Three more representative bands were recorded for Amx-HAp (Fig 1H) at 1776, 1612 and 1409 cm^−1^ that were assigned to C=O stretching of β-lactamic ring, aromatic C=C bending and symmetric COO^−^ stretching, respectively. These assignments confirmed that this antibacterial drug was physically mixed to HAp as previously reported (Bisson-Boutelliez). Col-HAp (Fig 1I) revealed additional FTIR absorption bands related to this structural protein. A signal recorded at 1655 cm^−1^ corresponding to amide I, where the peptide C=O groups contribute to stretching vibrations. Moreover, a band at 1545 cm^−1^ was assigned to the C–N stretching vibrations and N–H bending of amide II. An absorption band at 1245 cm^−1^ was related to amide III due to the stretching and binding vibrations of C–O and N–H, respectively.[14] Thus, collagen was also suitable blended to HAp as proposed.

### 3.2 Collection and characterization of human dental pulp-derived mesenchymal stem cells (hDPSCs)

Dental pulp samples of permanent teeth were obtained from three patients with a mean age of 27 years. Visual observation under bright field microscopy showed fibroblastroid morphology and capacity to adhere to plastic and, after isolation, took on average 20 days to proliferate to exceed 80% confluence (Fig 2A).

**Fig 2.**
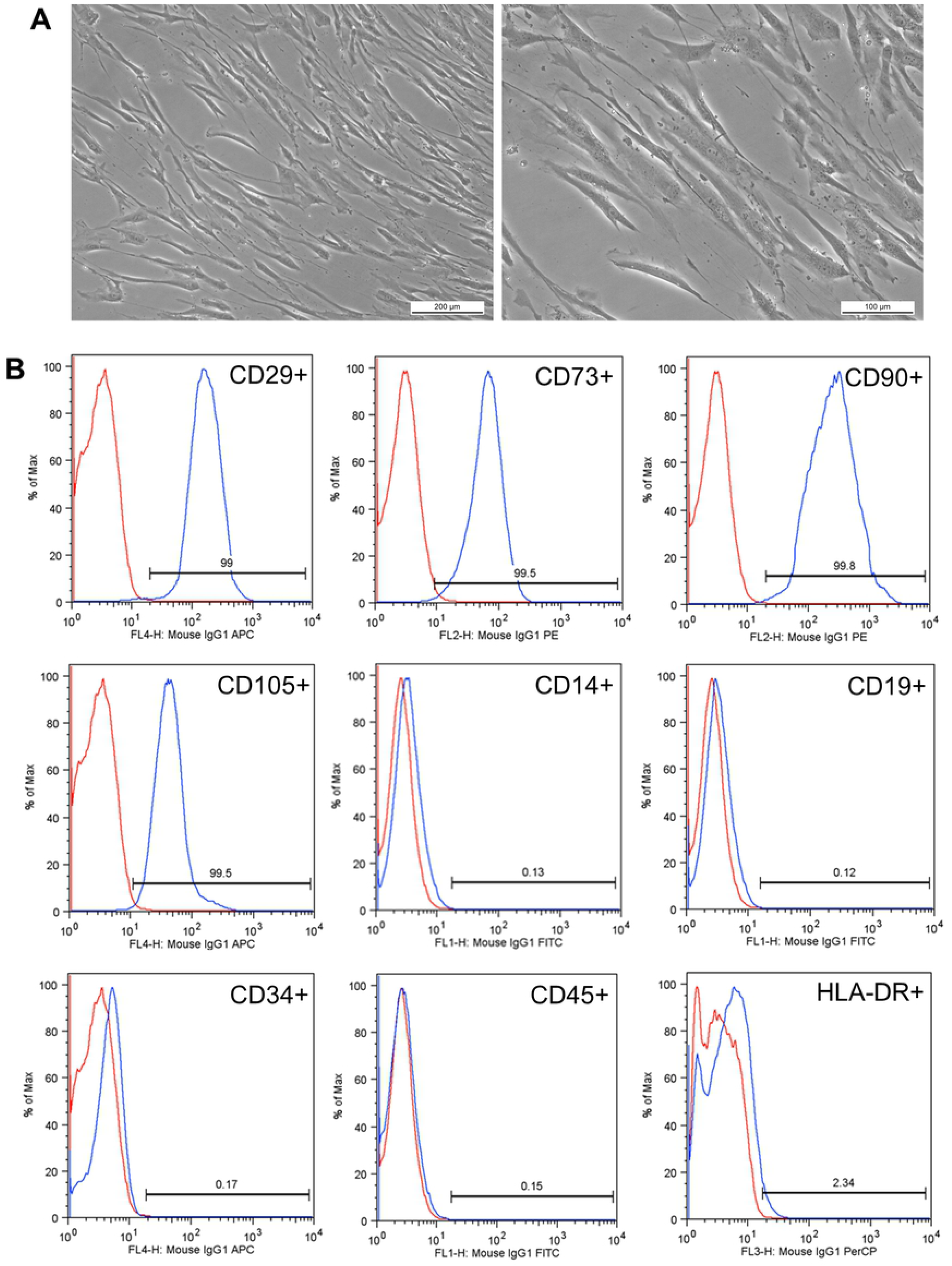
Characterization of hDPSCs. *A*. Representation of a sample in third passage. hDPSCs have plastic adherence and fibroblastoid morphology. Increases: 100x and 200x, respectively. *B*. hDPSCs surface markers of a study sample. In red, the isotype control is shown and in blue are represented the positive markers CD29, CD73, CD90, CD105, and negative markers CD14, CD19, CD34, CD45, and HLA-DR. Results expressed as a percentage (%).

#### 3.2.1 Characterization of human dental pulp-derived mesenchymal stem cells (hDPSCs)

The analysis of cell characterization showed an immunophenotypic profile compatible with the definition of mesenchymal stem cells as suggested by Dominici *et al*.[15] for all three samples. The mean scores for each marker were as follow: positive markers CD29 (98.97%), CD73 (99.17%), CD90 (99.63%), CD105 (99.27%), and negative for CD14 (0.22%), CD19 (0.09%), CD34 (0.12%), CD45 (0.21%), and HLA-DR (1.5%) (Fig 2B). The cells had a mean viability of 99.55% and 0.4% of the cells were Annexin-V stained which was indicative of apoptosis.

### 3.3 Osteogenic Differentiation

The ability of osteogenic differentiation was compared between the positive control and each biomaterial (HAp, Amx-HAp, and Col-HAp). The cells cultured with commercial osteogenic differentiation medium (positive control) demonstrated the presence of calcium crystals after Alizarin red S staining as well as those cultured with HAp-based materials in study (Fig 3).

**Fig 3.**
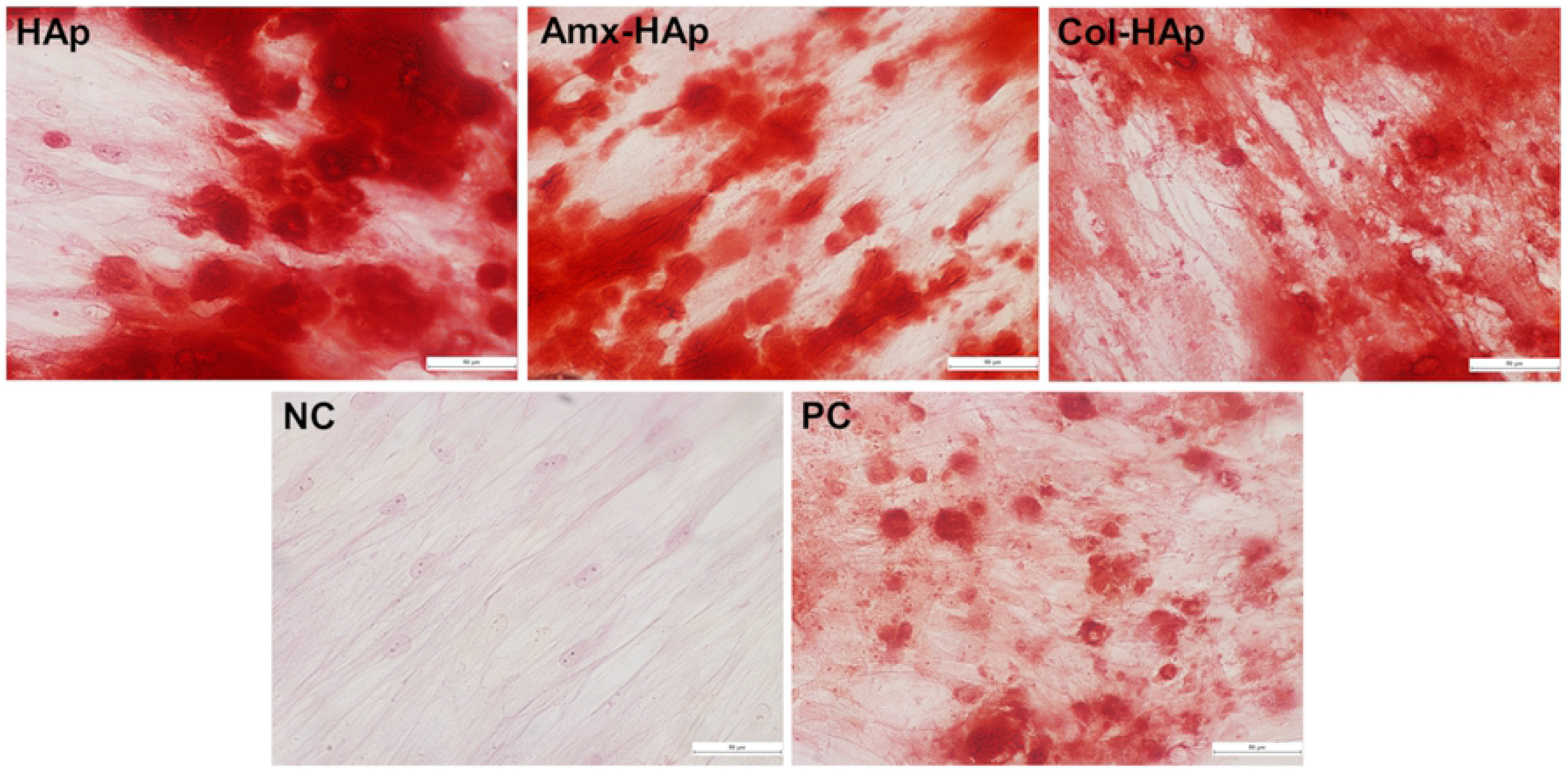
Induction of osteogenic differentiation of hDPSCs by the biomaterials tested in this study. HAp: hydroxyapatite nanosheet-assembled microspheres; Amx-HAp: amoxicillin-hydroxyapatite composite; Col-HAp: collagen-hydroxyapatite composite; NC: Negative control; PC: Positive control. The calcium crystals are present in groups HAp, Amx-HAp, Col-HAp, and positive control.

No difference in genes expression was observed after 21 days of induction. hDPSCs with no induction and those under different conditions of induction highly expressed collagen IA and osteonectin at mRNA, which indicated these genes played an important role in odontogenesis regardless of induction stimulus (Fig 4).

**Fig 4.**
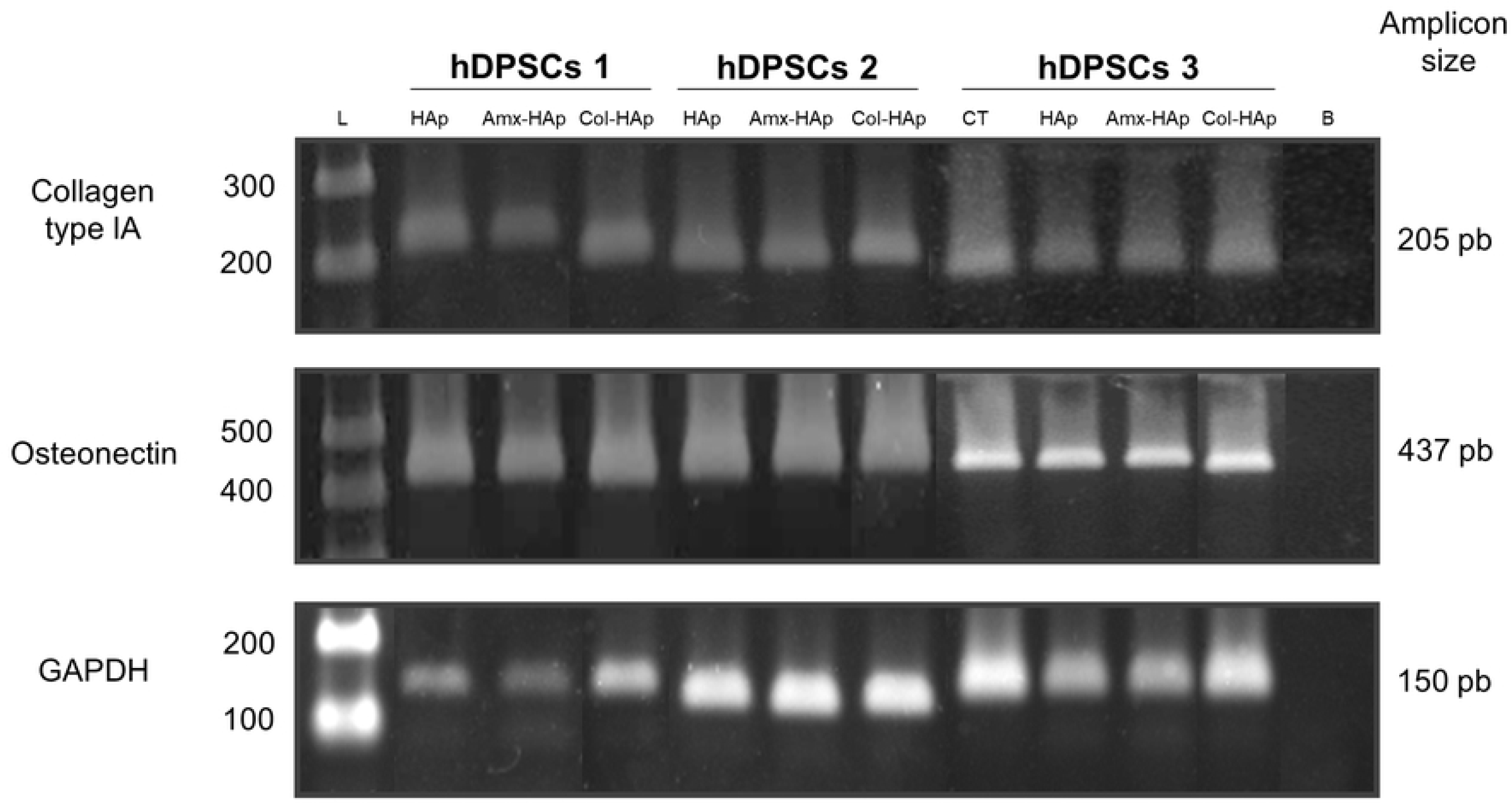
Expression of the collagen type IA and osteonectin markers of osteogenesis by PCR. Comparison between the control (hDPSCs with no osteogenic induction) and three samples of hDPSCSs (submitted to six different medium of osteogenic induction). CT: control; HAp: hydroxyapatite nanosheet-assembled microspheres; Amx-HAp: amoxicillin-hydroxyapatite composite; Col-HAp: collagen-hydroxyapatite composite; L: ladder, B: blank; GAPDH: glyceraldehyde-3-phosphate dehydrogenase, housekeeping gene; hDPSCs: dental pulp-derived mesenchymal stem cells.

### 3.4 Cytotoxicity Analysis

The cytotoxicity results from cell viability assay using 7-AAD dye showed that all groups had low mortality rate. The samples depicted the following mean ± SEM of stained non-viable cells: Control (1.473 ± 0.318%), HAp (1.417 ± 0.019%), Amx-HAp (1.403 ± 0.110%), and Col-HAp (0.757 ± 0.113%). Col-HAp group presented a number of non-viable cells statistically lower than the control group (*p*=0.03). The other groups did not demonstrate statistical difference when compared with the control group (p>0.05).

### 3.5 Migration Assay

The results obtained in the migration assay after 24 hours (Fig 5A) showed that the negative control was the only group with a statistically lower migration of hDPSCs when compared with the positive group (NC *vs*. PC, 0.3087 ± 0.2117 *vs*. 0.7840 ± 0.2493 nm, *p*=0.0008). In that sense, the biomaterials HAp, Amx-HAp, and Col-HAp revealed the same migration behavior for hDPSCs observed to the positive control. Col-HAp also provided a statistically significant higher migration of hDPSCs than HAp (Col-HAp *vs*. HAp, 0.9272 ± 0.3835 *vs*.0.5165 ± 0.1367 nm, *p*=0.0248).

**Fig 5.**
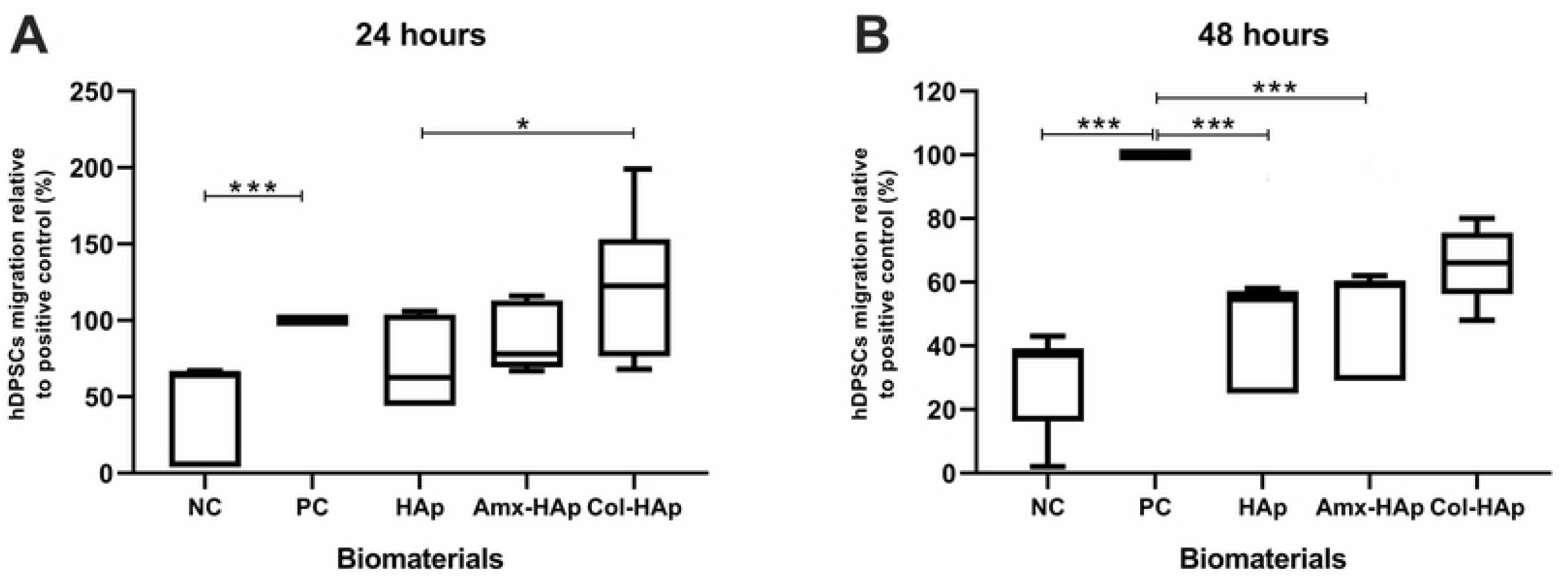
Migration assay of hDPSCs through 8-mm-pore membranes toward the tested biomaterials. *NC*: Control negative (IMDM with 1% FBS); *PC*: Control positive (IMDM with 20% FBS); HAp: hydroxyapatite microspheres; Amx-HAp: amoxicillin-hydroxyapatite composite; Col-HAp: collagen-hydroxyapatite composite. The positive control was normalized for 100% migration rate. Statistical analysis was performed using Kruskal-Wallis test with Dunn post-test; P-values: * <0.05, ** <0.01, *** <0.001.

In the 48-hour evaluation there was a significant difference between the negative control and the positive control (NC *vs*. PC, 0.3857 ± 0.1747 *vs*. 1.498 ± 0.6080 nm,*p* < 0.0001). The migration results obtained for the biomaterials HAp, Amx-HAp, and Col-HAp was intermediate from those achieved by control groups (Fig 5B). There was no statistical difference between positive control and Col-HAp (p>0.05).

## 4. Discussion

The studies concerning the synthesis of biomaterials such as hydroxyapatites with different particle sizes and forms are required to obtain suitable scaffolds in tissue engineering for achieving both osteoinductive and dentinogenic potential. Furthermore, advanced biomaterials inspired by drugs and biological molecules can broaden the prospects in terms of new technologies and variety of applications.[16]

In recent years, various HAp-based biomaterials with different morphologies such as nanowires [17], nanoneedles [18], or nanoflowers [19], and hierarchically nanostructured porous microspheres [20–23] or microflowers [24] have been synthesized. In addition, HAp are adequate candidates as drug or protein delivery carriers [25–27]. The biomaterials obtained showed crystalline peaks attributed to pure HAp. In that sense, both Amx and Col were incorporated as non-crystalline (amorphous) materials in composites structure.

The microwave-hydrothermal HAp and Amx-HAp demonstrated high superficial area since these biomaterials were arranged as nanosheet-assembled microspheres as previously reported.[28] These 3-D flower-like structures assembled with nanosheets radially oriented may impact the interaction with hDPSCs. Moreover, the Amx-HAp composite demonstrated more disordered and aggregated nanosheets than HAp, probably due to the β-lactamic antibiotic presence and the milling effect. The Col-HAp composite exhibited irregular microparticles with smooth surface, which resulted in a comparatively lower surface area. The FTIR spectra confirmed the main stretching and bending vibrations due to the main chemical groups of HAp, Amx, and Col. This analysis suggested that the composites were suitably obtained since no novel chemical bond were assigned during the biomaterials preparation.

After the hDPSCs immunophenotypic characterization, this study was focused on investigating if HAp, Amx-HAp, and Col-HAp may alter the induction of osteogenic differentiation of hDPSCs through Alizarin Red S staining and immunohistochemistry analysis since this analysis can reveal the presence of calcium stores in hDPSCs extracellular matrix.[29] The biomaterials HAp, Amx-HAp, and Col-HAp did not avoid the osteogenic differentiation of hDPSCs comparing to the control groups. Hence, these data evidence that these biomaterials *per se* are able to induce hDPSC osteogenic differentiation *in vitro*; nevertheless, the presence of osteogenic differentiation promoting factors accelerates and strengthens this process, as resulted from the substantial augmentation of number and size of calcium content in the extracellular matrix.[29] Moreover, it has been already shown that scaffolds containing HAp improved cell proliferation and promoted production of mineralized extracellular matrix more than that observed for the scaffold without HAp.[30]

Regarding to the cytotoxicity, no HAp-based biomaterial increased the percentage of non-viable cells when compared with the control group. Besides that, Col-HAp was able to preserve the cell viability significantly better than the control group. This is in line with data reported in literature, in fact, HAp appears to have great potential for bone tissue engineering as it showed no toxic effect on cell culture studies and also has good affinity of cellular attachment on the developed material surface. In addition, collagen is a natural component of bone tissue, where it stimulates mesenchymal stem cells to differentiate into osteoblasts, initiating new bone formation.[31] Collagen also increases water retention, which facilitates cell attachment.[32]

Considering that DPSCs are a vital source of osteoprogenitor cells that have a fundamental role in the mechanisms related to the pulp tissue plasticity,[33] the migration assay was performed. The biomaterials HAp, Amx-HAp, and Col-HAp showed a similar behavior than the positive control group in the migration assay after 24 hours. Col-HAp also demonstrated this result after 48 hours. Migration is a key property in the bone development and the treatment of bone defects since the cell chemotaxis and the invasion of cells through extracellular matrix are responsible to ensure the encouraging effect of a bone substitute biomaterial.[34,35] All the HAp-based biomaterials were feasible to stimulate hDPSCs migration. In this context, Col-HAp suitably acted as a chemoattractant agent due to its ability to provide a directional hDPSC response after 24 and 48 hours.

The reported evidences remarkable demonstrate that the osteo-inductive features of the studied HAp-based biomaterials, mainly Col-HAp scaffold, support the potential of hDPSCs inducing the formation of new bone tissue. These data suggest that these materials may represent a suitable tool to promote the migration, the proliferation, and the differentiation of bone cells by enhancing regeneration and fracture healing.

## 5. Conclusions

HAp, Amx-HAp, and Col-HAp were successfully obtained and characterized. These biomaterials proved to ensure the osteogenic differentiation of hDPSCs, to be non-cytotoxic and to stimulate hDPSC migration.

Col-HAp scaffold showed better features for the dynamic parameters of cell viability and cell migration capacities for hDPSCs. These data present high clinical importance because Col-HAp can be used in a wide variety of therapeutic areas, including ridge preservation, minor bone augmentation, and periodontal regeneration.

## Acknowledgments

This study was partially supported by Araucaria Foundation (Grant number 15/2017, Yasmine Mendes Pupo).

## Author Contributions

*Conceptualization*: Yasmine Mendes Pupo.

*Formal analysis*: Jessica Mendes Nadal.

*Investigation*: Yasmine Mendes Pupo.

*Methodology*: Lidiane Maria Boldrini Leite, Lisiane Antunes, Eliane Leal de Lara, Rafael Eiji Saito.

*Project administration*: Alexandra Cristina Senegaglia.

*Supervision*: Sandra Regina Masetto Antunes, Paulo Vitor Farago.

*Writing – original draft*: Yasmine Mendes Pupo.

*Writing – review & editing*: Yasmine Mendes Pupo, Paulo Vitor Farago.

